# The potential genetic network of human brain SARS-CoV-2 infection

**DOI:** 10.1101/2020.04.06.027318

**Authors:** Colline Lapina, Mathieu Rodic, Denis Peschanski, Salma Mesmoudi

## Abstract

The literature reports several symptoms of SARS-CoV-2 in humans such as fever, cough, fatigue, pneumonia, and headache. Furthermore, patients infected with similar strains (SARS-CoV and MERS-CoV) suffered testis, liver, or thyroid damage. Angiotensin-converting enzyme 2 (ACE2) serves as an entry point into cells for some strains of coronavirus (SARS-CoV, MERS-CoV, SARS-CoV-2). Our hypothesis was that as ACE2 is essential to the SARS-CoV-2 virus invasion, then brain regions where ACE2 is the most expressed are more likely to be disturbed by the infection. Thus, the expression of other genes which are also over-expressed in those damaged areas could be affected. We used mRNA expression levels data of genes provided by the Allen Human Brain Atlas (ABA), and computed spatial correlations with the LinkRbrain platform. Genes whose co-expression is spatially correlated to that of ACE2 were then clustered into 16 groups, depending on the organ in which they are the most expressed (as described by the NCBI genes database). The list of organs where genes sharing local over-expression with the ACE2 gene are the most expressed is astonishingly similar to the organs affected by Covid-19.

## Introduction

The coronavirus SARS-CoV-2 (previously named 2019-nCoV) first emerged in Wuhan, China, in late 2019. The disease then massively spread to Europe and other continents. It causes severe acute respiratory syndrome (SARS), and results in significant morbidity and mortality.

The most widely reported symptoms of SARS-CoV-2 in humans are fever, cough, fatigue, pneumonia, headache, hemoptysis, and dyspnea [1]. According to Huang et al. [2], 63% of patients also suffered from lymphopenia. All 41 patients developed pneumonia, with abnormal findings on chest CT. Complications included acute respiratory distress syndrome (29%), RNAaemia (15%), acute cardiac injury (12%), and secondary infection (10%). Furthermore, Chen et al. [3] included other symptoms like confusion (9%) and diarrhea (2% of patients).

This new form of coronavirus (SARS-CoV-2) shares a very high similarity with the severe acute respiratory syndrome CoV (SARS-CoV) and Middle East respiratory syndrome CoV (MERS-CoV), both in terms of RNA sequence and infection pathway [4,5].

Moreover, growing evidence reveals that coronaviruses can also invade the central nervous system (CNS), causing neurological symptoms. SARS-CoV infection is not always restricted to the respiratory tract, as it has been reported in the brains of patients and laboratory animals, where the brain stem was densely infected [6,7,8].

In 2002 and 2003, studies on SARS patients established the presence of SARS-CoV particles in the brain, found in both neurons and gliocytes [9,10,11]. Based on experimental studies on transgenic mice, Netland et al. and Li et al. [12,13] concluded that SARS-CoV and MERS-CoV is likely to enter the brain through olfactory nerves when administered intranasally. From there, viruses would promptly spread other areas of the brain, including thalamus and the brain stem. It should be noted that in mice infected with low inoculum doses of MERS-CoV, particles have been exclusively found in the brain, but not in the lungs. This indicates that the CNS infection was more decisive in the high mortality detected in infected mice [13].

It was reported that Angiotensin-converting enzyme 2 (ACE2) is the main receptor for SARS-CoV and MERS-Cov [12,14]. Recently, Baig et al. [15] demonstrated that SARS-CoV-2 also uses ACE2 to target the central nervous system (CNS).

As ACE2 membrane proteins are the entry point for several strains of coronavirus into human cells, we can assume that brain regions where the gene coding for ACE2 is expressed the most are also the regions that get the most infected by the virus. Therefore, biological functions handled by other genes expressed in these regions are likely to be affected by the infection.

To detect the most expressed genes in the regions where ACE2 is over expressed, we used ABA [15], the only atlas covering mRNA expressions in the whole human brain. All the mRNA expressions were integrated in LinkRbrain platform [17]. Based on the over-expression of these mRNA, LinkRbrain performed topographical correlations between the genes.

The aim of this study is to understand the potential impact of a SARS-CoV-2 infection in the brain. Hypotheses were drawn using the 20,789 mRNA expressions of the genetic base of Allen Institute for Brain [15], and on spatial correlations provided by the platform LinkRbrain [17], we identified the gene networks correlated to ACE2. Finally, we discovered a strong relationship between these gene networks and the symptoms registered in patients with coronavirus.

## Results and discussion

In this study we explored the topographical network of genes that are over-expressed in the same regions as ACE2.

Our first objective was to topographically localize the mRNA expressions of ACE2. Consequently, we explored the topographical network of genes with the highest mRNA expressions in the same brain regions.

Thanks to the topographical distance calculation [17], LinkRbrain computed correlations between regions where ACE2 has the highest expression levels, and the coordinates of the over-expressions of each of the 20,788 other genes. From this calculation, 100 genes in particular (from a model of 20,789 genes) showed a significant increase in concentration of their mRNAs in the same regions where ACE2 mRNA levels are the highest. NCBI and GenesCards platforms were used to explore the other organs where these 100 genes are most expressed.

All these results can be observed interactively on LinkRbrain platform via the link http://authorized2019.linkrbrain.org/query/#5937, or in the generated file referenced as supplementary material (SM) 1. The 100 most correlated genes to ACE2 are listed, and we found information on 75 genes about the organs where they are the most expressed (see the list in SM2).

Analyzing the biological functions of these 75 genes allowed us to categorize them into 16 groups (see Table 1). The first group concerns olfactory receptors (which can be the entry point for the virus into the brain), but also genes that are most expressed in the organs related to the symptoms reported in the literature, such as the lymph nodes (for immunity), testis, spleen, liver, and intestines. Ding et al. [9] reported that the coronavirus was detected in some organs like lungs or intestine, However, no trace of this virus was detected in damaged organs such asSARS-CoV-damaged organs like oesophagus, thyroid, spleen, lymph node, bone marrow, testis, ovary, uterus, or heart.

**Table 1:**
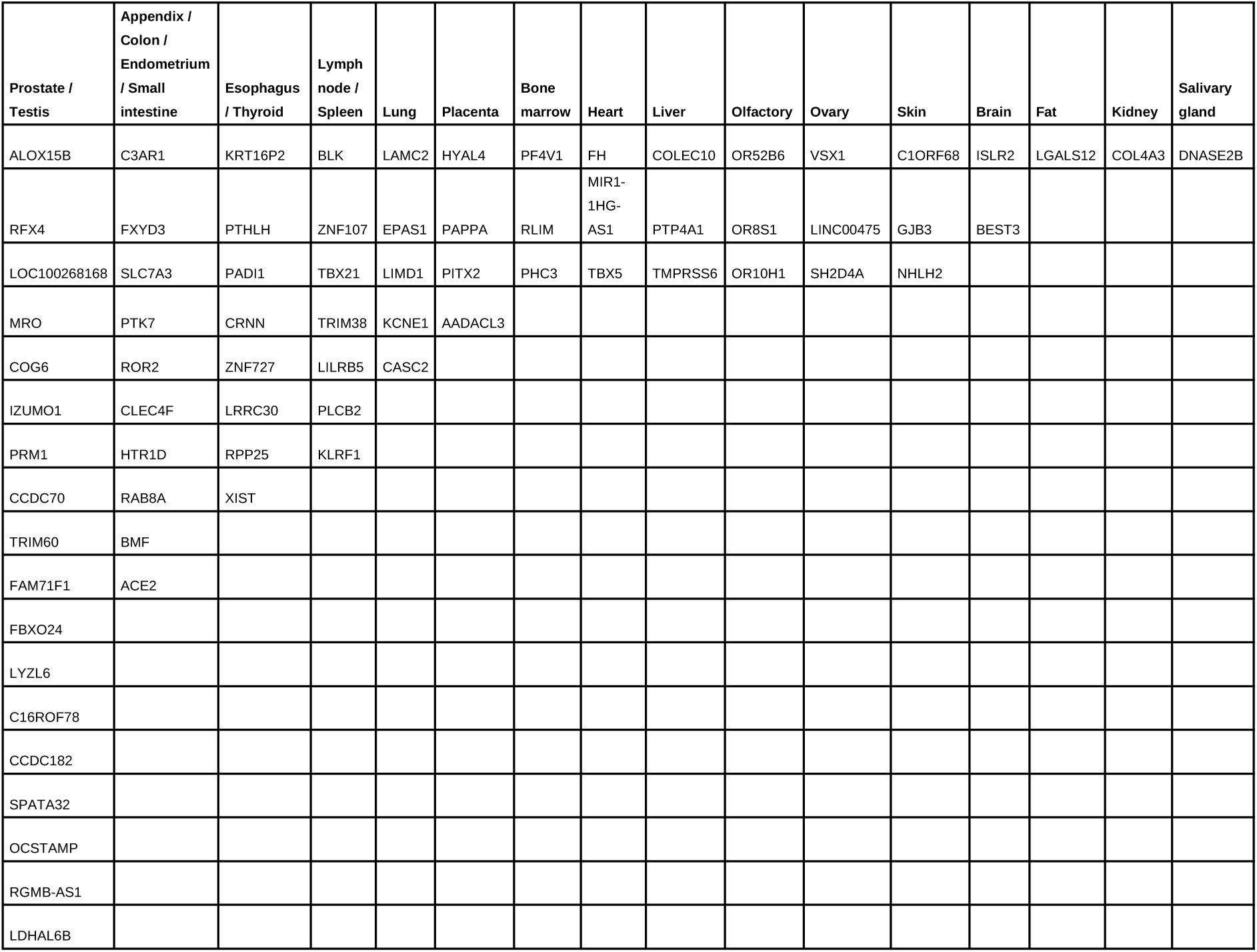
Genes co-expressed with ACE2 clustered by dominant biological function.

In summary, the list of organs where genes sharing local over-expression with the ACE2 gene are the most expressed is astonishingly similar to that of the organs affected by Covid-19.

### Olfactory group

By calculating correlations based on geometric distance, we identified three genes coding for olfactory receptors (OR10H1, OR52B6, OR8S1). Netland et al. [12] demonstrated that the brain of transgenic mice was a target organ for infection in SARS-CoV (human angiotensin-2 converting enzyme). Besides, they presented the olfactory bulb as the main entrance for the virus to reach the mouse brain. Our results showing genes corresponding to olfactory receptors in this topographical network are in agreement with these conclusions. Anosmia (loss of the ability to detect smells) has recently been identified as a distinctive symptom of the SARS-Cov-2 infection by several observations from clinicians and patients.

### Testis, prostate, and kidney

Another group includes eighteen genes (see the complete list in table 1) which play important roles in testes, kidneys, and the urinary tract, such as ALOX15B, LYZL6, CCDC70, and COL4A3 [18,19,20].

In several studies [2,3,21], SARS-CoV2 patients with abnormal kidney function or even kidney damage in addition to respiratory injuries have been mentioned, and the corresponding mechanism is unknown.

### Small intestine, appendix, colon, endometrium, and liver

In addition to common respiratory symptoms such as cough and fever, some patients may also experience other symptoms like diarrhea and liver damage [22,23], further challenging the patient’s recovery. Based on our topographical network of highly correlated mRNA expression with ACE2, we obtained a third category of 10 genes, which are strongly involved in the digestive system, such as BMF, C3AR1, FXYD3, and SLC7A3 (see the complete list in table 1) [24,25].

In the animal model, Yang et al. [26] presented the intestine as a target of SARS-CoV invasion. Furthermore, Xiao et al. [27] showed a human gastrointestinal infection of SARS-CoV-2.

### Lymph node, spleen, and bone marrow

One essential group detected within this topographical network is strongly linked to the immune system. Indeed, we have distinguished seven genes such as KLRF1, LILRB5, BKL and PHC3 (see the complete list in table 1), which are linked to the ACE2 topographical network and expressed in lymph node, spleen, and bone marrow [28,29,30,31].

In most SARS autopsies, both extensive necrosis of the spleen and atrophy of the white pulp with severe lymphocyte depletion have been found [32, 33]. Furthermore, several anomalies appeared in lymph nodes, such as atrophy [10] or necrotic inflammation [34].

### Thyroid gland

From the 75 studied genes, we found four genes with a mRNA expression highly geometrically correlated to the ACE2 mRNA expression that are expressed in the thyroid [35,36].

In their work Gu & Korteweg [37] reported, in the thyroid glands of five SARS autopsies, the destruction of epithelial cells with significant changes in the follicular architecture.

## Conclusion

Genes co-expressed with ACE2 in the brain reflect the pathogenic and even symptomatic picture of SARS-CoV-2 in an astonishing way. This could imply that when SARS-CoV-2 is linked to ACE2, the topographical network of genes overexpressed in the same regions are impacted.

Our results seem to be a serious hypothesis to explain the impact of SARS-CoV on affected patient’s organs like oesophagus, thyroid, spleen, lymph node, bone marrow, testis, ovary, uterus, and heart.

## Material and methods

We computed spatial correlations between the brain areas where mRNA of ACE2 is overexpressed, and the localized over expressions of 20,789 identified mRNA, using the genetic database provided by the Allen Institute for Brain [15]. We used ABA because it is the only atlas covering mRNA expressions in the whole human brain.

### Transcriptomic map provided by Allen Human Brain Atlas

ABA is a transcriptomic map of the whole human brain created by the Allen Institute [15]. RNAs were extracted from each brain to implement 58,693 complementary RNA hybridization probes, thus providing a RNA profile for the given subject.

The brain was then split into 946 samples, their coordinates being determined in MNI-space. The complementary RNA probes were used to assess the concentration of every fragment in each one of the samples, using RNA chips, thus providing the mRNA expression level of 20,789 genes in 946 known points of the MNI-space.

Each one of the 946 is by the concentration value of 20,789.

To ensure the reproducibility of the data obtained, two different brains were ensured [38].

These results have enabled a thorough study of the human genome expression at the cortical and sub-cortical levels.

### Analysis from LinkRbrain platform

LinkRbrain (www.linkrbrain.org) is an open-access web platform for multi-scale integration and visualization of knowledge on the brain. It integrates anatomical, functional, and genetic knowledge, provided by the scientific community. Genetic data were extracted from post-mortem studies, performed on brains from healthy subjects by the Allen Institute.

LinkRbrain allows the comparison of new results (set of coordinates) with previously published articles. Thus, the insertion of gene names in the platform allows their visualization as a new set of coordinates, although it establishes correlations with other genes whose level of expression is already identified at the same location.

The scoring algorithm described by Mesmoudi et al. [17] has been used to evaluate spatial correlations between the level of expression of ACE2 and each of the 20,789 genes present in the database.

In order to calculate the correlation between the inserted gene linked to the SARS-CoV, each set of coordinates, and the level of transcription relative to the 20,789 ABA genes present in LinkRbrain library, we used the platform internal topographical distance already defined in [17] (See the detail of this distance in SM3).

## Results viewing

Thanks to the visualization tools available on LinkRbrain we obtained:

- **2D and 3D visualizations**: these allow to envision the coordinates of the different transcription levels of ACE2 mRNA.
- **Topographical proximity graph:** this graph illustrates the 25 most closely correlated mRNA expressions.
- **List of genes:** based on the same scoring algorithm, LinkRbrain provides, among 20,789 mRNA gene expressions, a list of mRNAs which share the highest co-expression with the ACE2 mRNA.

### Functional genes study

We used NCBI (https://www.ncbi.nlm.nih.gov/gene/) [39] and GeneCards (www.genecards.org/) to conduct a functional study and the organ target of each considered gene. Indeed, we focused on the genes coding the mRNAs with the highest concentration at the coordinates where the highest expressions of the ACE2 gene were found.

## Acknowledgements

We would like to thank the Allen Institute for Brain Sciences for sharing their human brain genetic expression data. This study was funded by the French General Secretariat for investment (SGPI) via the National Research Agency (ANR) “Programme d’investissement d’Avenir (PIA ANR-10-EQPX-0021-01)” and by the CNRS innovation (prematuration program). The study was realized in the framework of the “équipement d’excellence MATRICE” headed by Denis Peschanski (matricememory.fr). This program is administratively sponsored by the HESAM Université.

## Author Contributions

Conceived and designed the experiments: SM. Performed the experiments: SM, MR, CL. Analyzed the data: SM, CL, MR. Wrote the paper: SM, CL, MR, DP.

## References

1. Adhikari SP, Meng S, Wu YJ, et al. Epidemiology, causes, clinical manifestation and diagnosis, prevention and control of coronavirus disease (COVID-19) during the early outbreak period: a scoping review. Infect Dis Poverty. 2020;9(1):29. Published 2020 Mar 17. doi: 10.1186/s40249-020-00646-x

2. Huang C, Wang Y, Li X, et al. Clinical features of patients infected with 2019 novel coronavirus in Wuhan, China [published correction appears in Lancet. 2020 Jan 30;:]. Lancet. 2020;395(10223):497–506. doi: 10.1016/S0140-6736(20)30183-5

3. Chen N, Zhou M, Dong X, et al. Epidemiological and clinical characteristics of 99 cases of 2019 novel coronavirus pneumonia in Wuhan, China: a descriptive study. Lancet. 2020;395(10223):507–513. doi: 10.1016/S0140-6736(20)30211-7

4. Yuan Y, Cao D, Zhang Y, et al. Cryo-EM structures of MERS-CoV and SARS-CoV spike glycoproteins reveal the dynamic receptor binding domains. Nat Commun. 2017;8:15092.

5. Wan Y, Shang J, Graham R, Baric RS, Li F. Receptor recognition by novel coronavirus from Wuhan: an analysis based on decade-long structural studies of SARS. J Virol. 2020. https://doi.org/10.1128/JVI.00127-20.

6. Li YC, Bai WZ, Hashikawa T. The neuroinvasive potential of SARS-CoV2 may play a role in the respiratory failure of COVID-19 patients [published online ahead of print, 2020 Feb 27]. J Med Virol. 2020;10.1002/jmv.25728. doi: 10.1002/jmv.25728

7. Bleau C, Filliol A, Samson M, Lamontagne L. Brain Invasion by Mouse Hepatitis Virus Depends on Impairment of Tight Junctions and Beta Interferon Production in Brain Microvascular Endothelial Cells. J Virol. 2015;89(19):9896–9908. doi: 10.1128/JVI.01501-15

8. McCray PB Jr, Pewe L, Wohlford-Lenane C, et al. Lethal infection of K18-hACE2 mice infected with severe acute respiratory syndrome coronavirus. J Virol. 2007;81:813–821.

9. Ding Y, He L, Zhang Q, et al. Organ distribution of severe acute respiratory syndrome (SARS) associated coronavirus (SARS-CoV) in SARS patients: implications for pathogenesis and virus transmission pathways. J Pathol. 2004;203:622–630. 32.

10. Gu J, Gong EC, Zhang B, Zheng J, Gao ZF, Zhong YF, Zou WZ, Zhan J, Wang SL, Xie ZG, Zhuang H, Wu BQ, Zhong HH, Shao HQ, Fang WG, Gao DX, Pei F, Li XW, He ZP, Xu DZ, Shi XY, Anderson VM, Leong ASY. Multiple organ infection and the pathogenesis of SARS. J Exp Med. 2005;202(3):415–424

11. Xu J, Zhong S, Liu J, et al. Detection of severe acute respiratory syndrome coronavirus in the brain: potential role of the chemokine mig in pathogenesis. Clin Infect Dis. 2005;41:1089–1096.

12. Netland J, Meyerholz DK, Moore S, Cassell M, Perlman S. Severe acute respiratory syndrome coronavirus infection causes neuronal death in the absence of encephalitis in mice transgenic for human ACE2. J Virol. 2008;82:7264–7275.

13. McCray PB Jr, Pewe L, Wohlford-Len Li K, Wohlford-Lenane C, Perlman S, et al. Middle East respiratory syndrome coronavirus causes multiple organ damage and lethal disease in mice transgenic for human dipeptidyl peptidase 4. J Infect Dis. 2016;213:712–722.

14. Hamming I, Timens W, Bulthuis ML, Lely AT, Navis G, van Goor H. Tissue distribution of ACE2 protein, the functional receptor for SARS coronavirus. A first step in understanding SARS pathogenesis. J Pathol. 2004;203(2):631–637. doi: 10.1002/path.1570

15. Baig AM, Khaleeq A, Ali U, and Syeda H. Evidence of the COVID-19 Virus Targeting the CNS: Tissue Distribution, Host–Virus Interaction, and Proposed Neurotropic Mechanisms. 2020. ACS Chemical Neuroscience Article ASAP, DOI: 10.1021/acschemneuro.0c00122

16. Hawrylycz MJ, Lein E.S, Guillozet-Bongaarts A.L., Shen E.H, et al. An anatomically comprehensive atlas of the adult human brain transcriptome. Nature 2012; 489:391–399

17. Mesmoudi S, Rodic M, Cioli C, Cointet JP, Yarkoni T, Burnod Y. LinkRbrain: multi-scale data integrator of the brain. J Neurosci Methods. 2015;241:44–52. doi: 10.1016/j.jneumeth.2014.12.008

18. Chen JB, Zheng WZ, Li YC, et al. (2016). Expression Characteristics of the Ccdc70 Gene in the Mouse Testis During Spermatogenesis. Zhonghua Nan Ke Xue. 2016;22(1):12–16.

19. Huang P, Li W, Yang Z, et al. LYZL6, an acidic, bacteriolytic, human sperm-related protein, plays a role in fertilization. PLoS One. 2017;12(2):e0171452. Published 2017 Feb 9. doi: 10.1371/journal.pone.0171452

20. Xia L, Cao Y, Guo Y, et al. A Novel Heterozygous Mutation of the COL4A3 Gene Causes a Peculiar Phenotype without Hematuria and Renal Function Impairment in a Chinese Family. Dis Markers. 2019;2019:8705989. Published 2019 Feb 10. doi: 10.1155/2019/8705989

21. Chan, J.F., et al., A familial cluster of pneumonia associated with the 2019 novel coronavirus indicating person-to-person transmission: a study of a family cluster. Lancet, 2020.

22. Chan HL, Kwan AC, To KF, Lai ST, Chan PK, Leung WK, Lee N, Wu A, Sung JJ. Clinical significance of hepatic derangement in severe acute respiratory syndrome. World J Gastroenterol, 11 (2005), pp. 2148–2153

23. Chai, X., et al., Specific ACE2 Expression in Cholangiocytes May Cause Liver Damage After 2019-nCoV Infection. bioRxiv, 2020: p. 2020.02.03.931766.

24. Hausmann M, Leucht K, Ploner C, et al. BCL-2 modifying factor (BMF) is a central regulator of anoikis in human intestinal epithelial cells. J Biol Chem. 2011;286(30):26533–26540. doi: 10.1074/jbc.M111.265322

25. Bibert S, Aebischer D, Desgranges F, et al. A link between FXYD3 (Mat-8)-mediated Na,K-ATPase regulation and differentiation of Caco-2 intestinal epithelial cells. Mol Biol Cell. 2009;20(4):1132–1140. doi: 10.1091/mbc.e08-10-0999

26. Yang XH, Deng W, Tong Z, et al. Mice transgenic for human angiotensin-converting enzyme 2 provide a model for SARS coronavirus infection. Comp Med. 2007;57(5):450–459.

27. Xiao F, Tang M, Zheng X, Liu Y, Li X, Shan H, Evidence for gastrointestinal infection of SARS-CoV-2 [published online March 3, 2020]. Gastroenterology. doi: 10.1053/j.gastro.2020.02.055

28. Roda-Navarro P, Hernanz-Falcón P, Arce I, Fernández-Ruiz E. Molecular characterization of two novel alternative spliced variants of the KLRF1 gene and subcellular distribution of KLRF1 isoforms. Biochim Biophys Acta. 2001;1520(2):141–146. doi: 10.1016/s0167-4781(01)00261-5

29. Hogan LE, Jones DC, Allen RL. Expression of the innate immune receptor LILRB5 on monocytes is associated with mycobacteria exposure. Sci Rep. 2016;6:21780. Published 2016 Feb 24. doi: 10.1038/srep21780

30. Crea F, Di Paolo A, Liu HH, et al. Polycomb genes are associated with response to imatinib in chronic myeloid leukemia. Epigenomics. 2015;7(5):757–765. doi: 10.2217/epi.15.35

31. Isono K, Fujimura Y, Shinga J, et al. Mammalian polyhomeotic homologues Phc2 and Phc1 act in synergy to mediate polycomb repression of Hox genes [published correction appears in Mol Cell Biol.2014 Jul;34(14):2771]. Mol Cell Biol. 2005;25(15):6694–6706. doi: 10.1128/MCB.25.15.6694-6706.2005

32. Wong RS, Wu A, To KF, Lee N, Lam CW, Wong CK, Chan PK, Ng MH, Yu LM, Hui DS, Tam JS, Cheng G, Sung JJ. Haematological manifestations in patients with severe acute respiratory syndrome: retrospective analysis. BMJ. 2003 Jun 21; 326(7403):1358–62.

33. Zhan J, Deng R, Tang J, Zhang B, Tang Y, Wang JK, Li F, Anderson VM, McNutt MA, Gu J. The spleen as a target in severe acute respiratory syndrome. FASEB J. 2006;20:2321–2328.

34. Lang ZW, Zhang LJ, Zhang SJ, Meng X, Li JQ, Song CZ, Sun L, Zhou YS, Dwyer DE. A clinicopathological study of three cases of severe acute respiratory syndrome (SARS). Pathology. 2003;35:526–531.

35. Xu Y, Wang J, Wang J. Long noncoding RNA *XIST* promotes proliferation and invasion by targeting miR-141 in papillary thyroid carcinoma. Onco Targets Ther. 2018;11:5035–5043. Published 2018 Aug 21. doi: 10.2147/OTT.S170439

36. Liu H, Deng H, Zhao Y, Li C, Liang Y. LncRNA XIST/miR-34a axis modulates the cell proliferation and tumor growth of thyroid cancer through MET-PI3K-AKT signaling. J Exp Clin Cancer Res. 2018;37(1):279. Published 2018 Nov 21. doi: 10.1186/s13046-018-0950-9

37. Gu, J, and Korteweg, C. Pathology and Pathogenesis of Severe Acute Respiratory Syndrome. Am J Pathol. 2007 Apr; 170(4): 1136–1147. doi: 10.2353/ajpath.2007.061088

38. Cioli C, Abdi H, Beaton D, Burnod Y, Mesmoudi S. Differences in human cortical gene expression match the temporal properties of large-scale functional networks. PLoS One. 2014;9(12):e115913. Published 2014 Dec 29. doi: 10.1371/journal.pone.0115913

39. Fagerberg L, Hallström BM, Oksvold P, et al. Analysis of the human tissue-specific expression by genome-wide integration of transcriptomics and antibody-based proteomics. Mol Cell Proteomics. 2014;13(2):397–406. doi: 10.1074/mcp.M113.035600

